# Optimal growth conditions and nutrient degradation characteristics of an aerobic denitrifying bacterium

**DOI:** 10.1101/353847

**Authors:** Sun Ling, Shi Juanjuan

**Affiliations:** School of Environmental Engineering, Xuzhou Institute of Technology, Xuzhou 221016, China

**Keywords:** aerobic denitrification, Arthrobacterium sp., isolate, bacterial identification, bacterial characteristics

## Abstract

An efficient aerobic denitrifying bacterium, QX1, identified as *Arthrobacter* sp. by morphological observations, physiological-biochemical tests, and 16s rDNA sequence analysis, was isolated from burdock fields. A phylogenetic tree for this strain was built based on the 16s rDNA sequence, and the effects of nitrogen source, carbon source, pH value, and temperature on the growth of the strain, in addition to its denitrification characteristics, were evaluated. The results showed that under conditions of a gas bath with shaking at 160 rpm, suitable growth conditions for this strain were LB medium plus 1% acetic acid sodium, 30°C, and pH 8. With these conditions, the strain grew to the logarithmic phase after about 4 h, the logarithmic phase was maintained for 16 h, steady growth was observed for 24 h, and growth began to decline thereafter. QX1 exhibited alkali resistance and could tolerate low (15°C) and high temperatures (40°C). At 30°C, pH 8, and shaking at 160 rpm, strains could degrade nitrate, with a removal rate of 89.92% within 48 h in synthetic sewage. Thus, strain QX1 could have applications as an alkaline-tolerant, broad-spectrum temperature-tolerant, efficient denitrifying bacterium in the field of denitrification repair.

## 1 Introduction

Traditional biological denitrification processes include autotrophic nitrification and anaerobic denitrification^[1–3]^. In recent years, heterotrophic nitrification and aerobic denitrification have become hot research topics in the field of biological denitrification. Several types of bacteria have been shown to function as aerobic denitrifying bacteria, including *Pseudomonas* sp.^[1,4,5]^, *Acinetobacter* sp.^[3,6,7]^, *Paracoccus* sp.^[2,8]^, *Rhizobium*^[9]^ *Alcaligenes* sp.^[10,11]^, *Bacillus* sp.^[12]^, and *Vibrio* sp.^[13]^. A strain of aerobic denitrifying bacteria, pseudomonas GL19, selected by Wang et al.^[14]^ showed resistance to low temperatures and could use a different nitrogen source for fast and efficient denitrification; this strain could also be applied for denitrification of wastewater and may have applications in wastewater denitrification treatment during winter. *Diaphorobacter* sp. PDB3^[15]^ has also been shown to remove ammonia nitrogen through cell assimilation and heterotrophic nitrification/aerobic denitrification, with a total organic carbon removal rate of up to 90.8%. The NO_3_− removal rate by *Pitt Acinetobacter* selected by Huang et al.^[16]^ reached 78.89% at 36 h under aerobic conditions with NO_3_− as the sole nitrogen source. Additionally, the NH_4_^+^, total nitrogen, and total organic carbon removal rates of *Pitt Acinetobacter* at 48 h were 95.25%, 80.42%, and 98.30%, respectively, when NH_4_^+^ was the only nitrogen source. Thus, biological denitrification technologies could be developed for use in practical applications.

In this study, we obtained a strain, designated QX1, from burdock planting field under aerobic conditions. We then aimed to analyze the morphological, physiological, and biochemical characteristics of this strain and to assess 16s rDNA sequences and denitrification characteristics in water to provide a theoretical basis for future applications of this strain in biological denitrification.

## 2 Materials and Methods

### 2.1 Bacterium source

The bacterium in this study was isolated from Xuzhou Venture burdock field obtained by the Xuzhou Engineering Institute of Environmental Engineering Laboratory Screening at 4°C.

### 2.2 Culture medium

The following media were prepared: 1) Enrichment medium, containing 2.0 g/L KNO_3_, 5.0 g/L C_6_H_12_O_6_, 1.0 g/L K_2_HPO_4_, 0.5 g/L KH_2_PO_4_, 0.2 g/L MgSO4·7H_2_O, 2 mL/L trace elements (pH 7.0-7.2). 2) LB medium, containing 5.0 g/L yeast extract, 10 g/L peptone, 5.0 g/L NaCl (pH 7.0-7.2; optionally, 25 g gelatin was added to make the medium solid). 3) BTB medium, containing 3.0 g/L beef extract, 10.0 g/L peptone, 15.0 g/L sucrose, 20.0 g/L NaCl, 0.01 g/L bromothymol blue (pH 7.0-7.2). 4) Denitrification medium, containing 5 g/L sodium citrate, 1 g/L KNO_3_, 1 g/L KH_2_PO_4_, 0. 5 g/L K_2_HPO_4_, 0.2 g/L Mg_2_SO_4_7H_2_O, 2 mL/L trace elements (pH 7.2). 5) BTB-denitrification medium, containing BTB medium plus denitrification medium at 4 mL/L (pH 7.0-7.2). 6) Selective medium for liquor, containing 5 g/L CH3COONa·3H_2_O, 0.05 g/L K_2_HPO_4_, 0.2 g/L KH_2_PO_4_, 0.2 g/L MgSO_4_·7H_2_O, 0.5 g/L CaCl_2_, 2 mL/L trace elements (pH 7.0-7.2). The trace element solution contained 1.5 g/L FeCl_3_·6H_2_O, 0.15 g/L H_3_BO_3_, 0.03 g/L CuSO_4_·5H_2_O, 0.03 g/L KI, 0.06 g/L Na_2_MoO4·2H_2_O, 0.12 g/L MnCl_2_·4H_2_O, 0.12 g/L ZnSO4·7H_2_O, and 0.12 g/L CoCl_2_·2H_2_O (pH 7.0).

### 2.3 Enrichment and selection of bacterial species

One gram of soil sample was collected in a 100-mL conical flask containing enrichment broth and shaken at 120 rpm at 26°C. The enrichment cycle was as follows: every 3 days, the bottle was stood for a moment for the precipitate to settle, after which 20 mL of the liquid was removed and fresh enrichment broth was added to a total volume of 100 mL. This method ensured that the denitrifying bacteria could continue to grow. Griess reagent was used to detect NO_2_-N, and diphenylamine-sulfuric acid reagent was used to detect the removal of NO_3_-N. When the solution turned colorless, the enrichment domestication process was considered complete.

Next, 0.5 mL bacteria after enrichment and domestication was removed and subjected to a gradient dilution (10^−1^–10^−9^). Twenty microliters of bacteria from the diluted samples was used to coat LB tablets, with three replicates for each dilution. Samples were maintained for 2-3 days in a biochemical incubator at 26°C. Mature colonies were transferred and streaked on plates; pure colonies were obtained after repeating this step 6-8 times. Pure colonies were inoculated in BTB-denitrification medium and developed for 2-3 days at 26°C; strains showing blue color were considered denitrifying bacteria. Selected denitrifying bacteria were cultured in 100 mL denitrifying medium at 26°C for 24 h with shaking at 160 rpm. The absorbance of NO_3_^−^-N and NO_2_-N was determined from the liquid after centrifugation. The denitrification rate was calculated according to the formula: η = (*A* – *A*_*t*_ – *B*_*t*_)/*A* × 100%, where · is the denitrification rate (%), *A* is the initial absorbance of NO_3_^−^-N, *A*_*t*_ is the absorbance of NO_3_^−^ in the solution after a certain time, *B*_*t*_ is the absorbance of NO_2_^−^ in the solution after a certain time. The strain with the highest denitrification rate was chosen for follow-up experiments.

### 2.4 Analysis of the morphological, physiological, and biochemical characteristics of the strain

Analysis of the morphological, physiological, and biochemical characteristics of the strain was carried out according to previously described methods^[17,18]^.

### 2.5 Amplification of the 16s rDNA gene by polymerase chain reaction (PCR)^[19]^ and construction of the phylogenetic tree

The 16s rDNA sequence was amplified from bacterial DNA using universal primers as follows: upstream (27FM) 5′-AGTTTGATCMTGGCTCAG-3′; downstream (1492), 5′-GGTTACCTTGTTACGACTT-3′. PCR was carried out using 50 ng/μL template (0.5 μL), 5× buffer (with Mg^2+^) 2.5 μL, 2.5 mM dNTPs (1 μL), 10 μM each primer (0.5 μL), 1 μM Taq-TM DNA polymerase (1 μL), with double distilled water to 25 μL. The PCR protocol included denaturation at 98°C for 3 min; 30 cycles of 98°C for 25 s, 55°C for 25 s, and 72 °C for 1 min; and a final extension at 72°C for 10 min. Agarose gel electrophoresis was used to analyze PCR products, and sequencing was performed by Sangon Biotechnology Co., Ltd. (Shanghai, China).

The 16s rDNA gene sequences were submitted to NCBI BLAST to retrieve similar sequences, and Clustal X1.83 software, based on the principles of maximum homology sequence alignment, with Mega5.0 software were used to implement the neighbor-joining method for construction of the phylogenetic tree. The stability of the branches was analyzed by the bootstrap method with 1000 repeats.

### 2.6 Effects of different conditions on growth

Fresh bacterial suspensions were inoculated into 250-mL conical flasks (150 mL) at an OD_600_ of 0.3. The cultures were then shaken at 160 rpm for 24 h, the OD_600_ was measured, and cultures were used for experiments below.

#### 2.6.1 Different nitrogen sources

Selective medium was added to mother liquor peptone (1%) and yeast extract (1%), with urea, NaNO_2_, KNO_3_, NaNO_2_, or KNO_3_ as the nitrogen source (molar ratio of C/N = 10). In addition, when the composite nitrogen source (LB) medium was prepared, the above six types of medium were used separately. Cultures were incubated with shaking at 26°C.

#### 2.6.2 Different carbon sources

The suitable nitrogen source was selected as described above. Then, sodium citrate CH_3_COONa·3H_2_O as the sole carbon source was replaced with 1% (v/v) sodium acetate, glucose, fructose, ethanol, potassium or sodium tartrate. This medium was then added to the selective medium in the mother liquor, and cultures were incubated at 26°C with shaking.

#### 2.6.3 Different temperatures

The optimal temperature was determined using the optimized sources of nitrogen and carbon. Temperatures of 15, 20, 25, 30, and 35°C were examined after incubation at 40°C.

#### 2.6.4 Different pH values

The optimal pH was determined by testing growth using the above optimized conditions at pH values of 5, 6, 7, 8, 9, and 10.

#### 2.6.5 Growth curves

A suitable amount of the bacterial suspension was selected to inoculate into 200 mL optimized culture medium. The culture was adjusted to an OD_600_ of 0.1 and incubated at the optimal temperature and pH. Samples were collected at certain times, and the absorbance at OD_600_ was determined to create bacterial growth curves.

### 2.7 Analysis of aerobic denitrification

A suitable amount of bacterial suspension was inoculated into 200 mL synthetic sewage (0.6 g/L glucose, 0.1 g/L peptone, 0.01 g/L yeast powder, 0.5 g/L sodium acetate, 0.05 g/L NaCl, 0.09 g/L KH_2_PO_4_·3H_2_O, 0.4g/L MgSO_4_·7H_2_O, 0.2 g/L KNO_3_). The OD_600_ was adjusted to 0.3, and bacteria were incubated with shaking under suitable conditions, with sampling every 4 h.

### 2.8 Analytical methods

Analyses were performed using the NO_3_^−^-N: phenol disulfonic acid method and NO_2_^−^-N:alpha naphthylamine spectrophotometry. Each experiment was repeated three times, and the results were averaged.

### 2.9 Data processing and statistics

Data were analyzed using Microsoft Excel software with single-factor analysis of variance and multiple comparisons. Differences with *P* values of less than 0.05 were considered statistically significant.

## 3 Results

### 3.1 Strain screening

From the selection procedures, 18 strains were initially picked and replated in LB culture medium. From these colonies, a total of seven single colonies were finally selected, named QX1-7. The screened strains were used for further analysis of denitrification rates, as shown in Table 1, and the strain QX1, which exhibited the highest denitrification rate, was used for further analyses.

**Table 1.** Strain denitrification rates under aerobic culture conditions after 24 h

| Project | Strain |  |  |  |  |  |  |  |
| --- | --- | --- | --- | --- | --- | --- | --- | --- |
|  | QX1 | QX2 | QX3 | QX4 | QX5 | QX6 | QX7 | QX8 |
| Denitrification rate (%) | 72.3 | 65.3 | 45.6 | 60.1 | 59.2 | 68.4 | 56.4 | 62.3 |

### 3.2 Morphological, physiological, and biochemical characteristics of QX1

QX1 colonies were pale yellow and neat, with a smooth, moist surface. For QX1, gram staining, transparent circle experiments, catalase assays, V-P experiments, nitrate reduction assays, and nitrate and oxidation origin gas assays were positive, whereas H_2_S analysis, semisolid AGAR piercing experiments, methyl red assays, and ammonia production were negative (Table 2).

**Table 2.** Physiological and biochemical characteristics of QX1

| Experiments | Phenomenon |
| --- | --- |
| Gram staining | — |
| Transparent circle experiment | + |
| Catalase experiment | + |
| Semi-solid AGAR puncture | — |
| V-P experiment | + |
| Methyl red experiment | — |
| Nitrate reduction experiments | + |
| Nitrate as a native gas | + |
| Ammonia production experiment | — |
| Oxidation of H<sub>2</sub>S | + |

“+” is positive, “—” is negative

### 3.3 16S rDNA gene PCR amplification and phylogenetic tree construction

The 16S rDNA sequence of strain QX1 was 1464 bp. An NCBI BLAST search showed that similarity was more than 99% compared with the sequences of *Arthrobacter nicotianae* SB42, *Arthrobacter arilaiti*, and *Arthrobacter nicotianae* and more than 98% compared with the sequences of *Arthrobacter arilaitensis, Arthrobacter* DSM12798, *Arthrobacter protophormiae* SSCT48, *Arthrobacter* sp. TVH. 003, and *Arthrobacter* sp. ZJUTW. The constructed phylogenetic tree is shown in Figure 1. Taken together, our data showed that this bacterium was within the genus *Arthrobacter*.

**Fig. 1.**
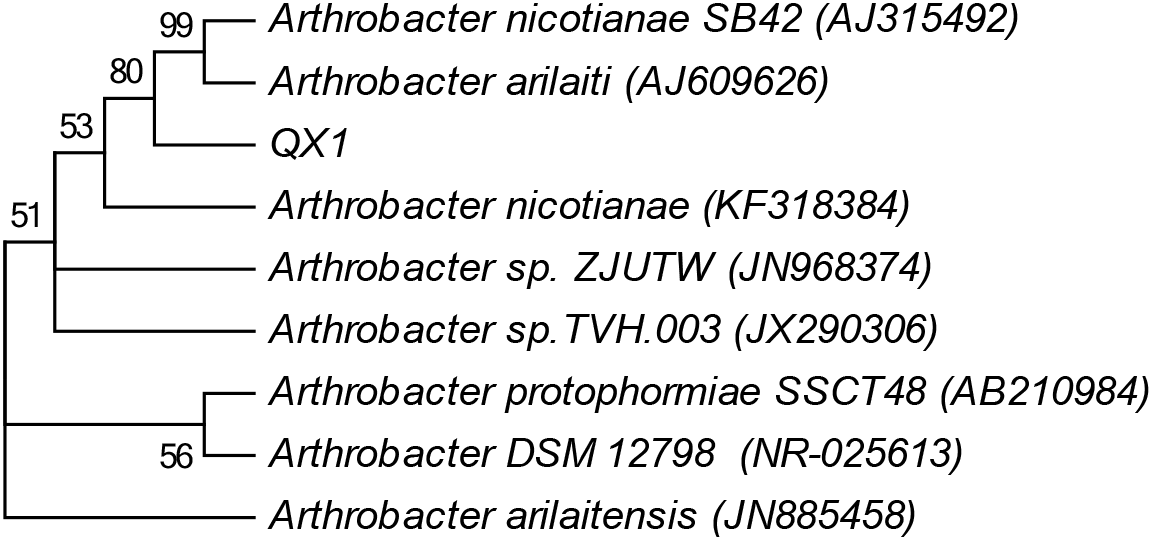
Phylogenetic tree of QX1 based on analysis of the 16S rDNA gene.

### 3.4 Effects of different conditions on the growth of QX1

#### 3.4.1 Effects of nitrogen source on growth

The optimum culture medium for strain QX1 was LB culture medium (Fig. 2a), with the following order of growth based on the nitrogen source: LB medium > peptone > yeast extract > urea > KNO_3_ > NaNO_2_ (OD_600_ values of 1.30, 1.24, 1.21, 1.13, 1.06, and 0.57, respectively). Strains grew better in the presence of organic nitrogen than in the presence of inorganic nitrogen, with highest growth seen for the composite organic nitrogen. Moreover, the differences in growth with different types of nitrogen sources were significant, with growth in LB medium significantly better than that in yeast extract medium (*P* < 0.05) or the three other types of culture medium (*P* < 0.01). Therefore, LB medium was used for subsequent analyses.

**Fig. 2.**
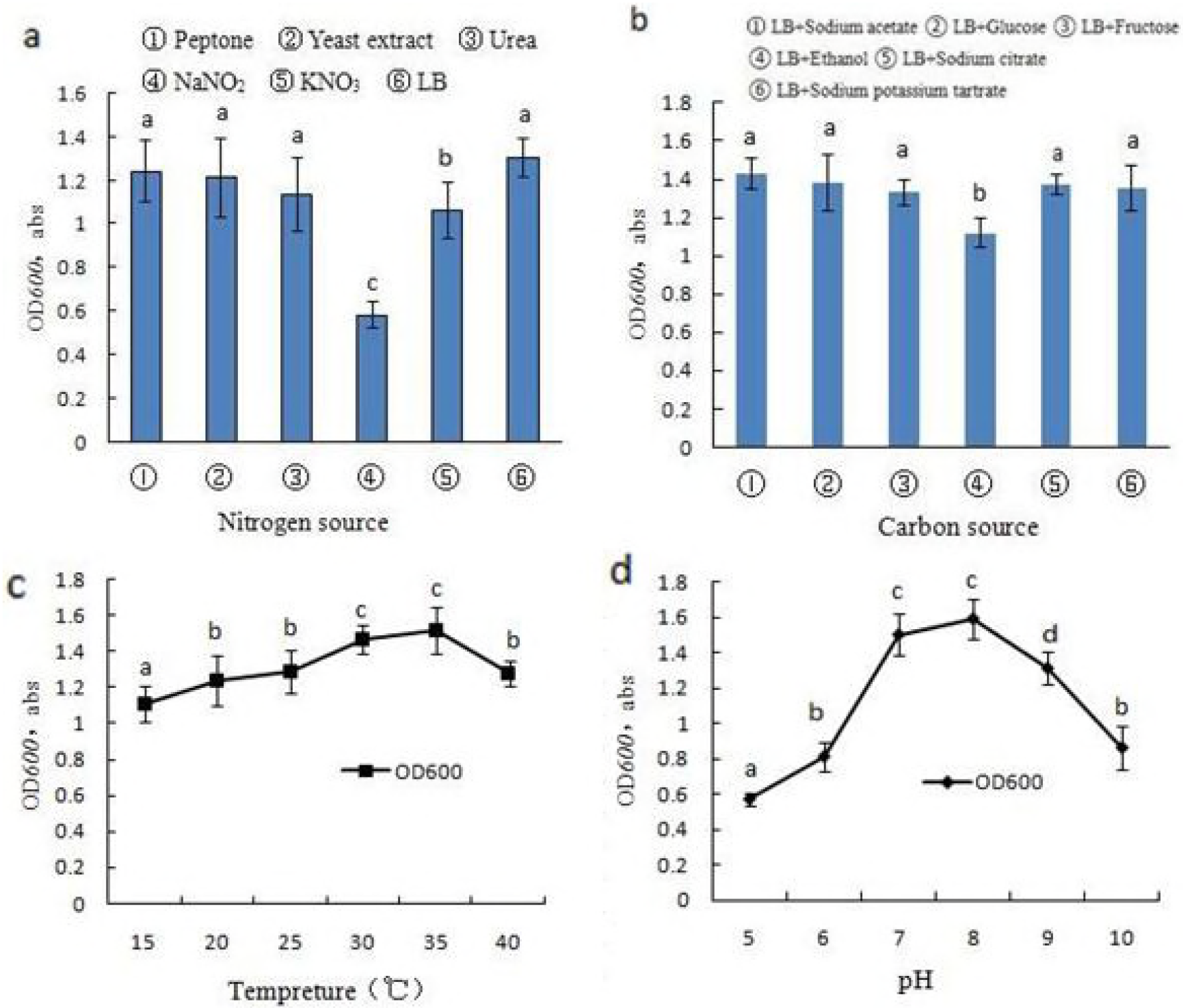
Effects of external conditions on the growth of QX1. Different letters represent significant differences, *P* < 0.05 or *P* < 0.01.

#### 3.4.2 Effects of carbon source on growth

The optimum carbon source was sodium acetate, with growth under conditions of different the carbon sources following the order of sodium acetate > glucose > sodium citrate > sodium potassium tartrate > fructose > ethanol (OD_600_ values of 1.43, 1.38, 1.37, 1.35, 1.33, and 1.12) after 24 h (Fig. 2b). Growth was significantly lower when ethanol was the carbon source compared with that of the other carbon sources (*P* < 0.05). Therefore, sodium acetate was used as the carbon source for subsequent analyses.

#### 3.4.3 Effects of temperature on carbon source

The growth of QX1 increased with increasing temperature from 15 to 35°C (OD_600_ values of 1.11, 1.24, 1.29, 1.47, and 1.52, respectively). However, at a temperature of 40°C, the OD600 decreased to 1.28 after 24 h (Fig. 2c). Thus, temperature affected the growth of QX1, and growth at 30 and 35°C was significantly better than that at other temperatures (*P* < 0.05 or *P* < 0.01, respectively). Therefore, we chose a temperature of 30°C for further experiments.

#### 3.4.4 Effects of pH value

As shown in Fig. 2d, pH affected the growth of QX1 as well. Optimal growth was observed at a pH of 8. Reduced growth was observed at lower pH, whereas strong growth was observed when the pH was in the range of 6-9. Statistical analysis showed that growth was significantly better at pH 7 and 8 than that at other pH values (*P* < 0.05). Moreover, growth was significantly reduced when the pH was 5 (*P* < 0.01) or when the pH was 6 or 10 (*P* < 0.05), indicating that strains could grow in a weakly acidic, neutral, or alkaline environment.

#### 3.4.5 Growth curve

Growth curves were created for QX1 based on the optimal culture conditions determined in the previous experiments (Fig. 3). Cells grew slowly during the adaptation phase from 0 to 4 h. However, after 4 h, QX1 cells grew quickly and entered the logarithmic phase, which lasted for approximately 16 h. After 20 h, the growth rate plateaued, exhibiting stable growth at 24 h. During this period of stable growth, the OD_600_ value was approximately 1.5–1.8, with bacterial counts of 1.756×10^13^−3.018 × 10^13^ cfu/mL. After 44 h, growth began to decrease. These differences in growth over time were significant.

**Fig. 3.**
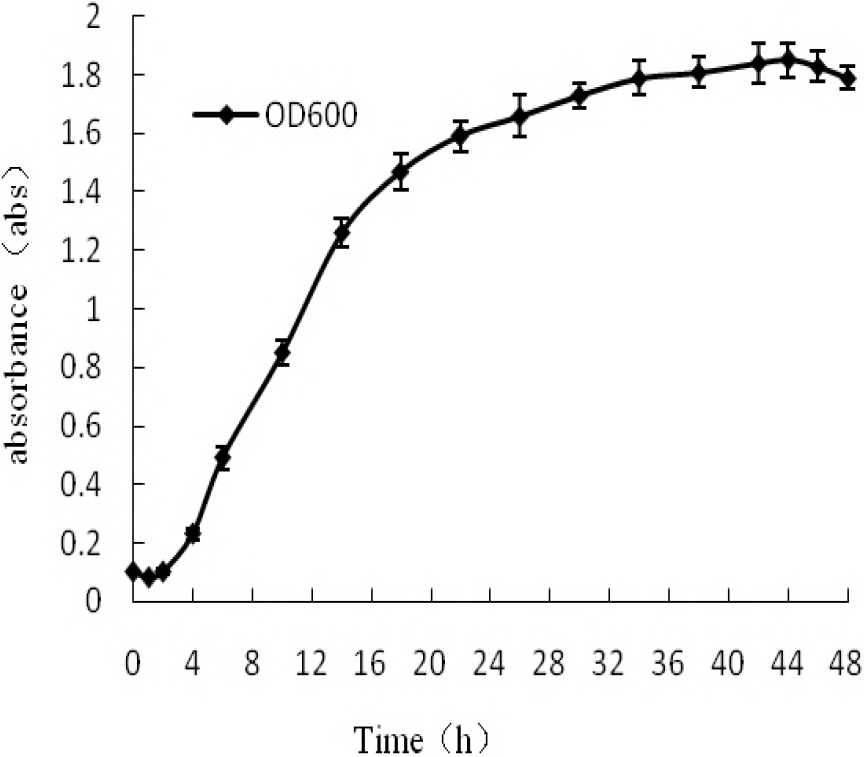
Growth curve for strain QX1

### 3.5 Denitrification performance of strain QX1

QX1 was inoculated into 200 mL synthetic sewage and incubated with shaking at 8,160 rpm at 30°C for 48 h, with 4-h sampling intervals. Nitrate nitrogen, nitrate nitrogen concentration changes, and nitrate nitrogen removal rates are shown in Fig. 4. The nitrate nitrogen content was reduced gradually from 200 to 20.36 mg/L. The maximum removal rate was 9.18 mg/L (20-24 h; 89.92%). Nitrite nitrogen did not accumulate, and levels remained low (0-0.06 mg/L). No ammonia nitrogen was detected throughout the entire process, similar to physiological and biochemical identification when the ammonia reaction was negative. Thus, the nitrate nitrogen gas also originated from air products. As shown in Fig. 4, QX1 was involved in denitrification mainly during the logarithmic phase of bacterial growth. This period is when bacterial growth and reproduction are most active; nutrition, energy, and cell synthesis ability are highest during this period, supporting that denitrification occurs primarily during this period.

**Fig. 4.**
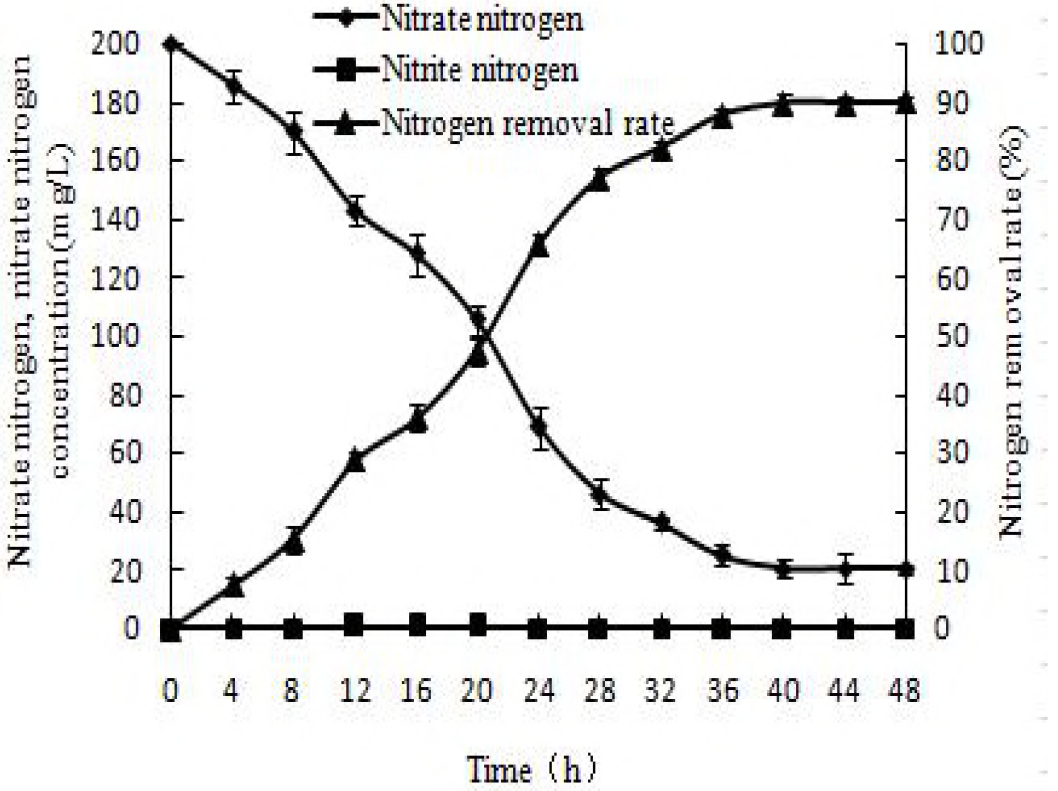
QX1 denitrification performance

## 4 Discussion

Denitrifying bacteria are primarily gram-negative bacteria, and most of the denitrifying gram-negative bacteria have been shown to belong to *Pseudomonas* sp., *Acinetobacter* sp., *Paracoccus* sp., *Alcaligenes* sp., and *Bacillus* sp. Here, we screened the denitrifying bacteria QX1 and identified this strain as belonging to the *Arthrobacter* genus through morphological observations, physiological-biochemical identification, and 16s rDNA sequence alignment. Generally, culture with gas bath shaking at a speed of 160 rpm is thought to yield cultures with saturated dissolved oxygen (DO) at 79-95%^[1]^. Additionally, under these conditions, the nutrient solution DO remained above 2 mg/L, according to actual measurements, and strains in BTB-denitrification medium showed blue halos, indicating that the bacteria had aerobic denitrification ability. Thus, we concluded these were aerobic denitrifying bacteria.

By selection and optimization of pH, temperature, nitrogen source, and carbon source, we determined the optimal growth conditions for strain QX1 as follows: LB medium containing 1% acetic acid sodium (mass-to-volume ratio), 30°C, and pH 8. QX1 aerobic denitrification occurred mainly during the logarithmic phase, consistent with the results of Wang^[20]^ and Fan^[21]^. There was no nitrite nitrogen accumulation during the process of QX1 denitrification, and the denitrification rate was 89.92% within 48 h. Aerobic denitrifying bacteria can also produce energy by pathways of aerobic denitrification and oxygen as an electron acceptor in the respiratory chain^[22]^. However, the production efficiency of the former is lower than that of the latter^[23]^, and with the abundance of DO, bacteria may prefer oxidation rather than aerobic denitrification. At the same time, the presence of a sufficient carbon source can result in the production of greater amounts of carbon skeleton, which is conducive to cell growth. In this study, we found that the strain QX1 could grow better in a composite organic nitrogen source and carbon source. Moreover, the QX1 degradation rate varies with the quality of the culture medium; when the bacteria exhibit a greater matrix absorption rate, the cells are in the “feed” state, and the bacteria do not expend their own nutrients and energy, whereas when the matrix absorption rate is slower, cells are in the “hunger” state and utilize their own nutrients and energy to maintain basal metabolism. Therefore, cells can only proliferate after surviving the “hunger” period^[24]^. This can explain the effects of different culture media on the OD600 values.

Temperature and pH can affect the growth of microbial and enzymatic reactions, and each bacterium has an optimum growth temperature and pH range. In this study, temperature and pH had a significant influence on QX1 growth. The optimum temperature was 30°C, and the optimum pH was 8; these values were similar to those of general microbes. At 40°C, the OD600 of QX1 was 1.28, whereas that at 15°C was 1.11, indicating that QX1 was resistant to low and high temperatures. Thus, this bacteria may be used as an alternative bacteria owing to its resistance to alkaline conditions and low or high temperatures, allowing efficient denitrification and supporting its broad potential applications.

## 5 Conclusions

In this study, an efficient aerobic denitrifying bacterium, QX1, was isolated from burdock fields. The strain QX1 was ultimately determined to be *Arthrobacter* sp. through morphological observations, physiological-biochemical identifications, and 16s rDNA sequence analysis. The optimal growth conditions for this strain were as follows: LB medium plus 1% acetic acid sodium, 30°C, and pH 8. This strain was resistant to alkaline conditions and to low and high temperatures. Finally, under optimal conditions, the nitrate nitrogen removal rate reached 89.92%. Thus, this bacteria may be used as an alternative bacteria for denitrification owing to its resistance to alkaline conditions and low or high temperatures, supporting its broad potential applications.

## Acknowledgements

The study was supported by grants from the Projects of Ministry of Housing and Urban-Rural Development under Grant No. 2016-R2-013 and the Young Project of Xuzhou Institute of Technology under Grant No. XKY2016122.

